# Ser/Thr kinase Trc controls neurite outgrowth in *Drosophila* by modulating microtubule-microtubule sliding

**DOI:** 10.1101/780999

**Authors:** Rosalind Norkett, Urko del Castillo, Wen Lu, Vladimir I. Gelfand

**Affiliations:** Department of Cell and Developmental Biology, Feinberg School of Medicine, Northwestern University, Chicago, IL 60611, USA

## Abstract

Correct neuronal development requires tailored neurite outgrowth. Neurite outgrowth is driven by microtubule sliding – the transport of microtubules along each other. We have recently demonstrated that a “mitotic” kinesin-6 (Pavarotti in *Drosophila*) effectively inhibits microtubule-sliding and neurite outgrowth. However, mechanisms of Pavarotti regulation in interphase cells and specifically in neurite outgrowth are unknown. Here, we use a combination of live imaging and biochemical methods to show that the inhibition of microtubule sliding by Pavarotti is controlled by phosphorylation. We identify the Ser/Thr NDR kinase Tricornered (Trc) as a Pavarotti-dependent regulator of microtubule sliding in neurons. Further, we show that Trc-mediated phosphorylation of Pavarotti promotes its interaction with 14-3-3 proteins. 14-3-3 binding is necessary for Pavarotti to interact with microtubules and inhibit sliding. Thus, we propose a pathway by which microtubule sliding can be up or down regulated in neurons to control neurite outgrowth, and establish parallels between microtubule sliding in mitosis and post-mitotic neurons.

## Introduction

In order to communicate, neurons must develop an extensive and precisely regulated network of axons and dendrites, collectively called neurites. Studying the mechanisms that form these processes is key to understanding early nervous system development. Neurites are filled with cytoskeletal components including microtubules. Neurons are exceptionally dependent on microtubules for long range transport of cargo. Also, microtubule organization is essential for powering initial neurite outgrowth (Kapitein and Hoogenraad, 2015; Lu et al., 2013; Winding et al., 2016). In order to drive initial neurite outgrowth, microtubules themselves become the cargo and are transported relative to each other by molecular motors – a process known as microtubule sliding. Indeed, in cultured *Drosophila* neurons, microtubules can be seen pushing the plasma membrane at the tips of growing processes (del Castillo et al., 2015; Lu et al., 2013). Previous work from our group has identified the classical kinesin – Kinesin-1 – as the motor responsible for the majority of microtubule sliding in neurons (Lu et al., 2013; Winding et al., 2016).

Observation of microtubule sliding in neurons is of particular interest as this process is best described during vast cytoskeletal reorganization in mitosis, rather than in terminally differentiated neurons (Baas, 1999). Microtubule sliding is observed in young neurons in culture, but decreases as neurons mature (Lu et al., 2013). Therefore, in addition to promoting neurite extension via microtubule sliding, there must also exist mechanisms to downregulate this process. This prevents overextension of neurites when their intended synaptic targets are correctly reached. Work from our group and others has previously identified the kinesin-6 Pavarotti/MKLP1 as a powerful inhibitor of microtubule-microtubule sliding. Depletion of Pavarotti/MKLP1 by RNAi leads to axon hyperextension and more motile microtubules (Del Castillo et al., 2015; Lin et al., 2012). Identifying a neuronal role for this kinesin was of interest as Kinesin-6 has well studied roles in mitosis. It exists as a heterotetramer with MgcRacGAP (Tumbleweed in Drosophila) to form the centralspindlin complex (Adams et al., 1998; Basant and Glotzer, 2018; Mishima et al., 2002). This complex bundles microtubules at the bipolar spindle in late anaphase (Hutterer et al., 2009). Here, it can locally activate RhoA and promote assembly of the contractile actin ring at the cortex, and so, cytokinesis (Basant and Glotzer, 2018; Verma and Maresca, 2019).

How Pavarotti itself is temporally regulated to inhibit microtubule sliding as neurons mature is unknown. Mitosis exhibits tight temporal regulation. We speculated that similar mechanisms might be at play in regulating Pavarotti activity in neurons with regard to microtubule sliding. One well studied facet of centralspindlin regulation in regards to mitotic progression is that of phosphorylation (Douglas et al., 2010; Guse et al., 2005). Based on bioinformatics and a literature search, we targeted Ser/Thr kinases known to modify Pavarotti during mitosis and tested their ability to modulate microtubule sliding in interphase cells. One potential kinase was the NDR kinase Tricornered (Trc, LATS in mammals) – shown to phosphorylate MKLP1 at S710 (the human ortholog of Pavarotti, S745) *in vitro* (Okamoto et al., 2015) (Fig 1 A). Trc regulates cell cycle exit (Hergovich et al., 2006) and also has conserved roles in neurite outgrowth, described in *Drosophila*, *C. elegans* and mammals (Emoto et al., 2006, 2004; Gallegos and Bargmann, 2004; Ultanir et al., 2012). How this kinase acts warrants further investigation.

**Figure 1.**
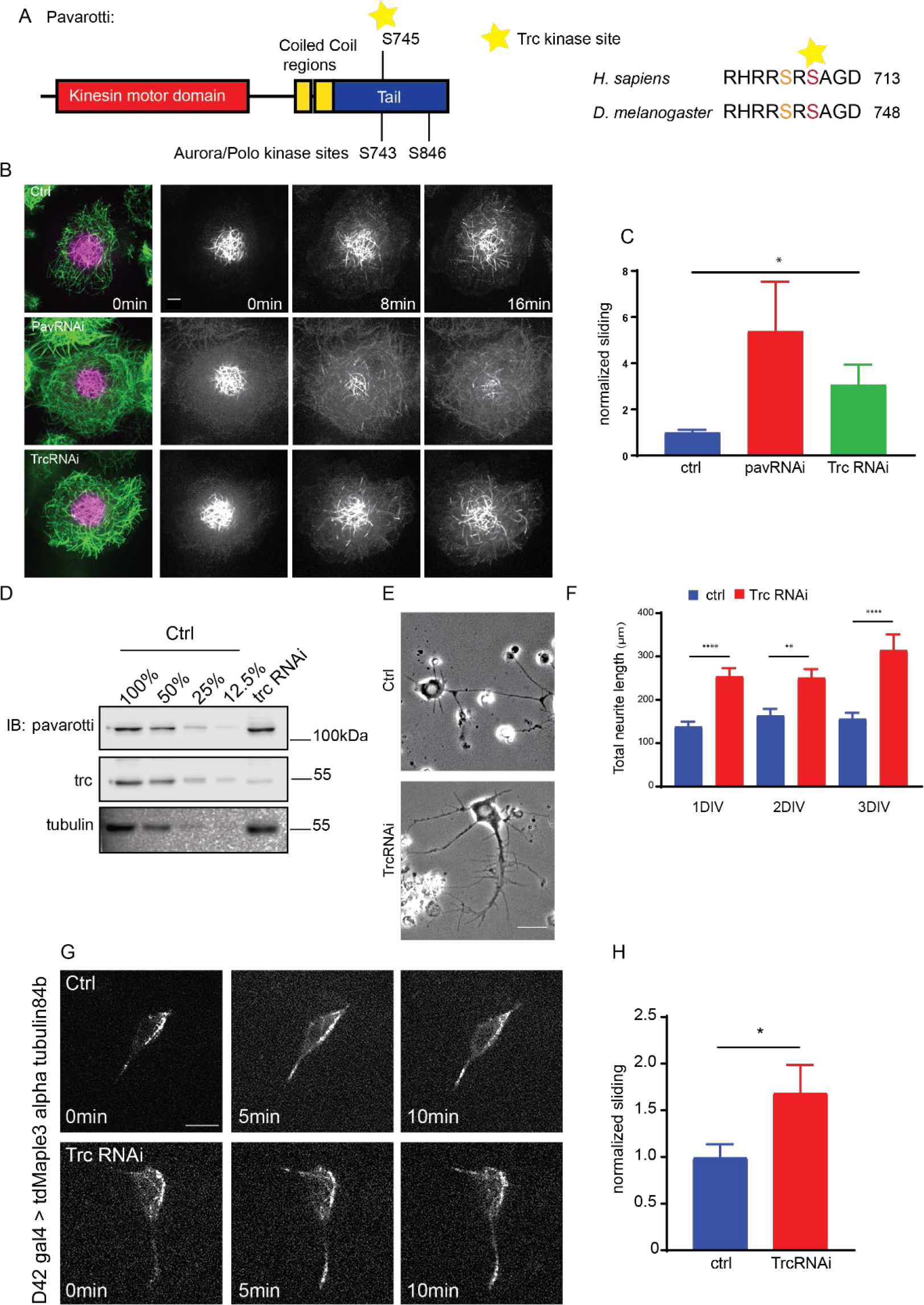
The kinase Trc regulates neurite outgrowth and microtubule sliding. A. Schematic showing domain structure and location of proposed Trc phosphorylation site in *Drosophila* Pavarotti and Human MKLP1. B. Example images of timelapse imaging to measure microtubule sliding in *Drosophila* S2 cells. C. Quantification of microtubule sliding rate shows an increase upon Pavarotti or Trc depletion. D. Western Blot of S2 cell extract showing efficient Trc knockdown. Pavarotti levels are unaffected. E. Representative images of 3^rd^ instar larvae cultured neurons under control or elav>Trc RNAi conditions. F. Quantification of total neurite length per cell over time in culture. The total neurite length is increased from control upon Trc depletion. 24hrs; ctrl = 137.8 ± 12.0μm, Trc RNAi 254 ± 19.1μm. 48hrs; ctrl = 163.7 ± 15.3μm, Trc RNAi = 251.2 ± 19.9μm, 72hrs; ctrl = 156.3 ± 13.8 μm, TrcRNAi = 314.4 ± 36.5μm. N = 11-23 cells from 3 independent experiments G. Example images from timelapse imaging of photoconverted microtubules in neurons under control conditions or upon Trc depletion. Tubulin was labelled with tdMaple3 alpha tubulin84b. After photoconversion, cells were imaged every minute for 10minutes. Scale bar = 5μm H. Quantification of microtubule sliding rates. Trc depletion leads to an increase in microtubule sliding rates in neurons. Ctrl = 1.0 ± 0.14, TrcRNAi = 1.69 ± 0.30. N = 20 cells from 3 independent experiments. P = 0.04 Student’s T-test

Here we use *Drosophila* S2 cells, neuronal culture and *in vivo* imaging to show Trc regulates microtubule sliding and dendrite outgrowth in neurons. We validate Pavarotti as a Trc substrate and demonstrate that phosphorylation of Pavarotti at S745 by Trc is necessary for proper control of microtubule sliding. We also show that phosphorylation of Pavarotti affects its subcellular distribution via interaction with 14-3-3 proteins in interphase cells – a mechanism conserved from mitosis. We demonstrate the function of this pathway in regulating development of *Drosophila* neurons.

## Results

### Tricornered Kinase inhibits neurite outgrowth and microtubule sliding

We have previously demonstrated the requirement of microtubule-microtubule sliding, by kinesin-1, for neurite outgrowth in *Drosophila* (Lu et al., 2013; Winding et al., 2016). This sliding is opposed by the mitotic kinesin-6 ‘Pavarotti’/MKLP1 (Del Castillo et al., 2015). However, the mechanism by which Pavarotti itself is regulated in this neuronal context is unclear. We hypothesized that Pavarotti may be regulated by phosphorylation, as in mitosis (Basant and Glotzer, 2017; Guse et al., 2005)(Fig 1 A). We targeted kinases known to modify Pavarotti during mitosis (AuroraB, Plk1 and Trc), and tested their ability to modulate neuronal development and microtubule sliding in non-dividing cells. Based on preliminary experiments we chose to focus on the NDR kinase Trc. Initially, we measured sliding using the model system of S2 cells, a *Drosophila* cell line. We have previously demonstrated that kinesin-1 carried out microtubule-microtubule sliding in these cells and that Pavarotti inhibits this (Del Castillo et al., 2015; Jolly et al., 2010). We decreased Trc levels using dsRNA. As shown in Fig 1 D, Trc is depleted and Pavarotti levels are unaffected. To measure microtubule sliding, we expressed a photoconvertible tubulin probe (tdEos-αTubulin84c) in S2 cells. Tubulin was photoconverted in a region of interest, and this specific population of microtubules was imaged by timelapse confocal microscopy. Sliding is measured as percentage of photoconverted tubulin outside the original photoconversion zone – see Methods. We found a significant increase in microtubule sliding upon depletion of Trc compared to control. (Fig 1 B, quantified in C, video 1).

Having established that Trc regulates microtubule sliding in S2 cells, we were next interested in investigating the potential role of Trc in neurite development. We analyzed primary neuronal cultures from third instar larvae expressing Trc RNAi (neuron specific expression was achieved by elav gal4>UAS Trc RNAi). We found a dramatic increase in total neurite length per cell *in vitro* (Fig 1 E and F).

Next, we directly tested the ability of Trc to regulate microtubule sliding in *Drosophila* cultured neurons. To do this, we expressed the photoconvertible maple tubulin under the control of the motor neuron specific D42 Gal4 driver and prepared dissociated neuronal cultures from brains of 3^rd^ instar larvae (Fig 1 G, video 2). Photoconversion was carried out in a constrained region of the cell and photoconverted signal was imaged over time to determine microtubule sliding rate, in a similar fashion to S2 cells. We compared control cells to those expressing Trc RNAi under the same driver. Consistent with our data in S2 cells, we found that depleting Trc levels led to an increase in microtubule sliding rate in primary culture (Fig 1 H). Therefore, Trc has the ability to modulate microtubule sliding in order to control neuronal neurite outgrowth.

Together, these data describe a role for Trc as a negative regulator of neuronal development, independent from cell division, and suggest that the mechanism by which Trc regulates neurite outgrowth is via microtubule sliding. This effect could be intrinsic, rather than dependent on external cues, as the effect can be seen in dissociated cultures.

### Trc kinase activity is necessary to control microtubule sliding

We also confirmed that this effect on sliding was dependent upon the kinase activity of Trc in two ways. Firstly, we depleted Furry (Fry) in S2 cells. Furry is a large protein with no known functional domain, shown to promote Trc kinase activity without affecting expression level (Fig 2 B) (Emoto et al., 2006). Upon knockdown of Fry, and so decrease in Trc kinase activity, we found an increase in microtubule sliding (Fig 2 B, C, video 3). Secondly, we depleted the endogenous Trc with a dsRNA targeting a non-coding region and expressed either WT Trc, constitutively active Trc (Trc CA), or kinase-dead Trc (Fig 2 D, E, video 4) (He et al., 2005). We confirmed expression of each of these mutants by fluorescence microscopy for BFP-Trc. We found that while WT and constitutively active Trc were able to reduce microtubule sliding to similar levels as control samples, the kinase dead mutant was not. Therefore, Trc negatively regulates sliding in a kinase dependent manner.

**Figure 2.**
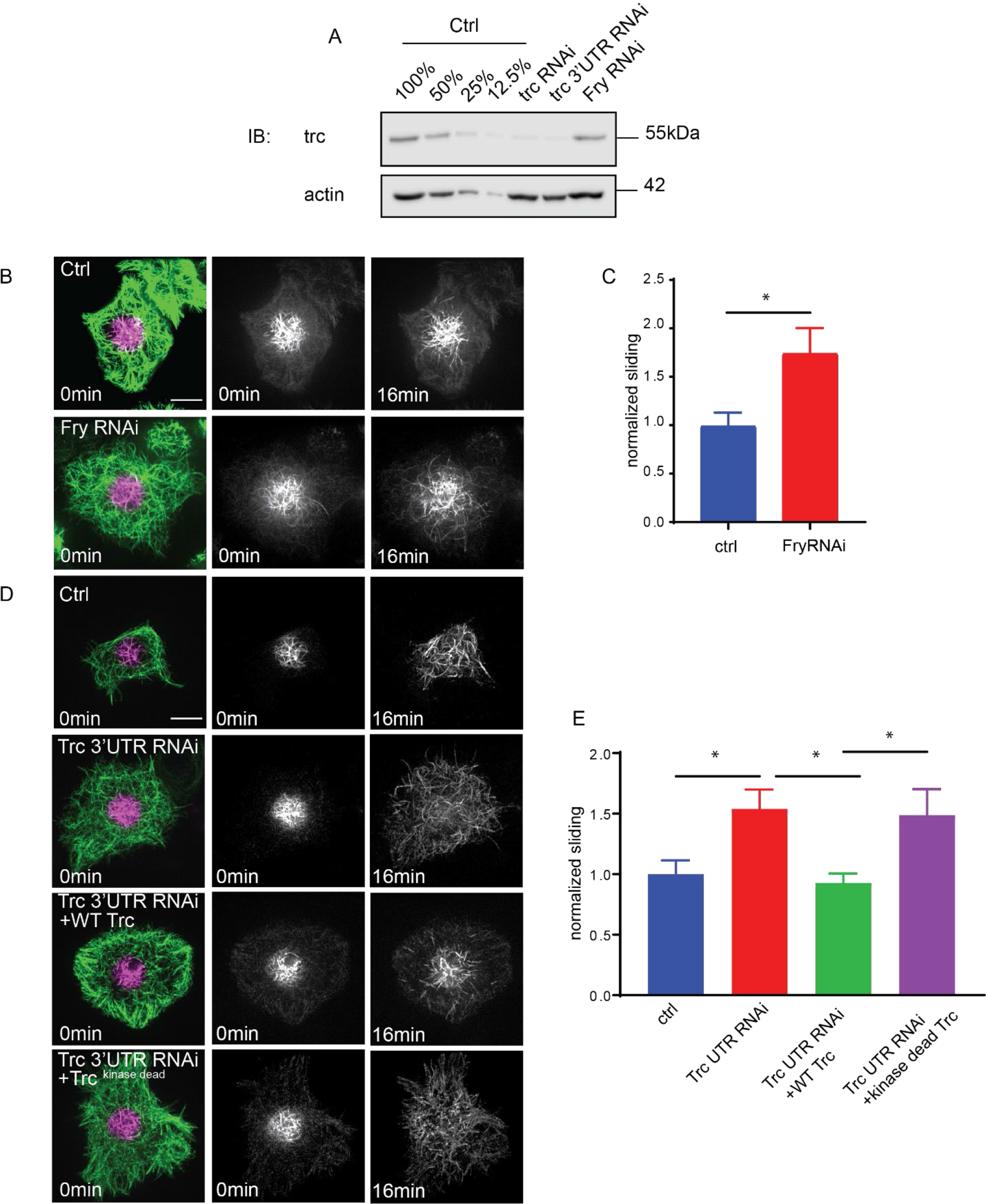
Trc regulates microtubule sliding in kinase dependent manner. A. Western Blot of S2 cell extract showing efficient Trc knockdown with a dsRNA targeting the Trc 3’UTR and dsRNA targeting Fry. Fry knockdown does not affect Trc protein level. Scale bar = 5μm. B. Sliding experiments in S2 cells show increased sliding upon depletion of Fry as quantified in C. n=26-37 cells from 4 independent experiments. Ctrl = 1 ± 0.13, Fry RNAi = 1.75 ± 0.26. Lower 95% CI ctrl = 0.72, Fry RNAi 1.23. Upper 95% CI ctrl = 1.268, Fry RNAi = 2.265. p= 0.024 Student’s T-test. D. Sliding experiments in S2 cells show increased sliding upon treatment with Trc 3’UTR dsRNA. This is rescued to control levels with expression of exogenous WT Trc, but not kinase dead Trc. Quantified in E. Scale bar = 5μm. n=46-50 cells from 4 independent experiments. Ctrl=1.0 ± 0.11, Lower 95% CI 0.77, Upper 95% CI 1.23, Trc3’UTR RNAi=1.54 ± 0.16, 1.218, 1.86, Trc3’UTR RNAi + WT Trc = 0.93 ± 0.08, 0.77, 1.09 Trc3’UTR RNAi + kinase dead Trc = 1.49 ± 0.22, 1.05, 1.92. Ctrl vs Trc 3’UTR RNAi p = 0.03, Trc 3’UTR vs 3’UTR + WT p = 0.01, 3’UTR + WT vs 3’UTR kinase dead p = 0.03. One-way Anova with Sidak’s post hoc correction.

### Tricornered Kinase phosphorylates Pavarotti to brake microtubule sliding

Next, we sought to investigate if this effect of Trc was dependent upon Pavarotti. In order to address this, we carried out sliding assays in S2 cells. We overexpressed Trc, either alone, or in conjunction with Pavarotti knockdown. Overexpression of Trc in S2 cells resulted in a decrease in microtubule sliding. This is in good agreement with our previous data describing Trc as a negative regulator of sliding (Fig 3 A, B, video 5). Importantly, upon depletion of Pavarotti, this decrease in sliding was lost (Fig 3 A, B). Therefore, Pavarotti must be present for Trc to oppose microtubule-microtubule sliding. In order to confirm Pavarotti was indeed phosphorylated by Trc at the predicted site S745, we carried out immunoblotting with a phospho-specific antibody for Pavarotti S745. We expressed GFP Pavarotti in HEK 293T cells and performed pull downs with anti GFP antibody. Western blot analysis showed basal phosphorylation at S745. Mutation of Pavarotti Ser745 to Ala gave no signal with the phospho-specific antibody, confirming the antibody specificity. Importantly, ectopic expression of constitutively active Trc (Trc CA), resulted in a roughly twofold increase in Pavarotti phosphorylated at S745. Therefore, Trc phosphorylates Pavarotti at Serine 745 in cells (Fig 3 I).

**Figure 3.**
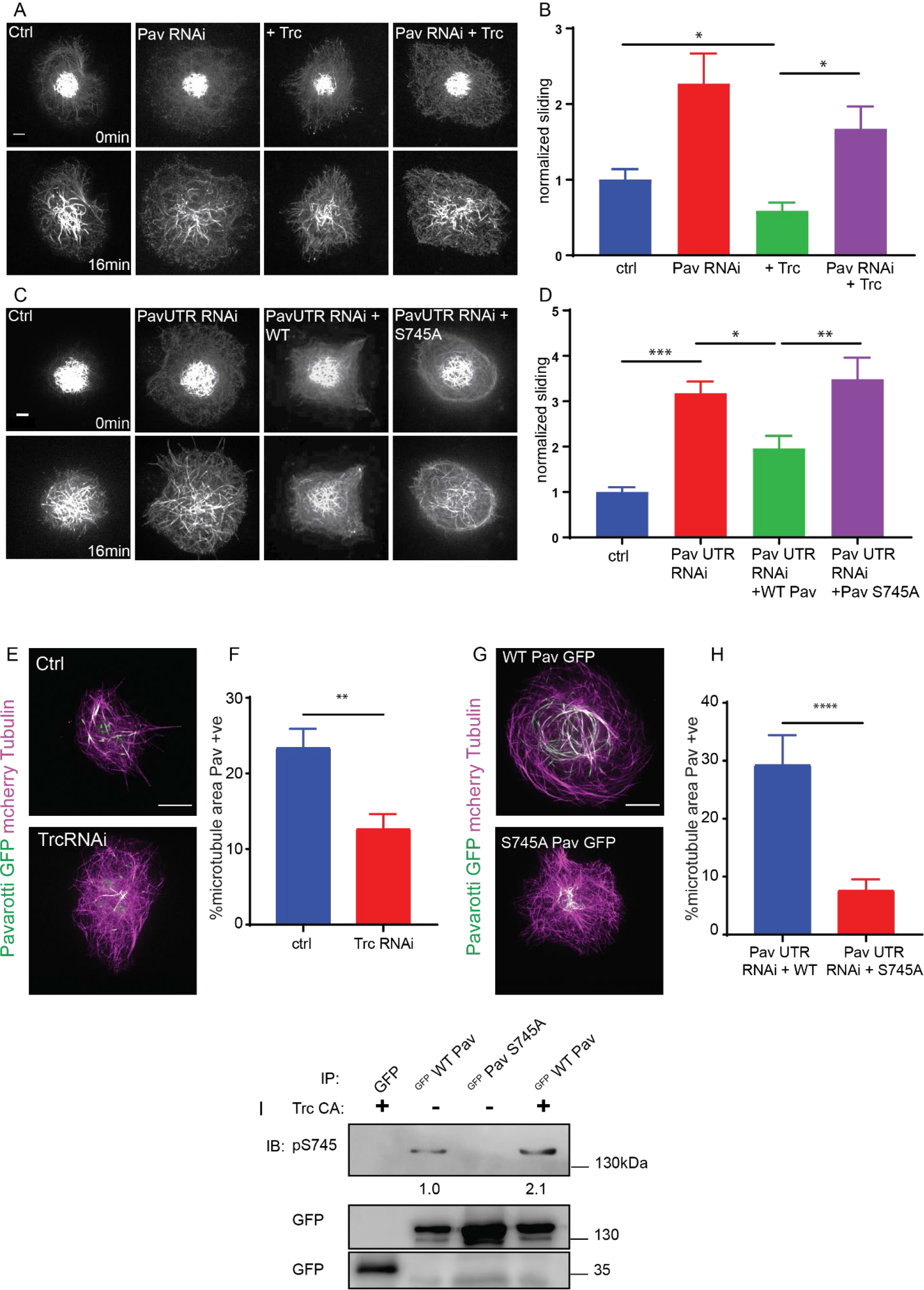
Trc regulates microtubule sliding via phosphorylation of Pavarotti. Sliding experiments in S2 cells show a decrease in microtubule sliding with Trc overexpression. Trc overexpression in conjunction with depletion of Pavarotti increases sliding beyond control levels as quantified in B. Scale bar 5μm C. Sliding experiments in S2 cells show an increase in microtubule sliding with Pavarotti depletion. The effect can be rescued with WT Pavarotti but not with Phospho null mutant S745A as quantified in D. n=39-48 cells from 4 independent experiments. Ctrl = 1 ± 0.10 Upper 95% CI = 1.2, Lower 95% CI = 0.79, Pav RNAi = 3.16 ± 0.26, 2.65, 3.70, Pav RNAi + WT = 1.96 ± 0.28, 1.40, 2.52 Pav RNAi + Pav S745A = 3.49 ± 0.48, 2.53, 4.44. Ctrl vs Pav RNAi p = 0.0001, Pav RNAi vs pav RNAi + WT p = 0.04, Pav RNAi vs Pav RNAi + S745A p = 0.94, Pav RNAi + WT vs Pav RNAi + S745A p = 0.0025. One-way ANOVA with Sidak’s post hoc correction. E. Example images of extracted S2 cells expressing mCherry Tubulin and WT Pavarotti GFP under control or Trc RNAi conditions. F Quantification of microtubule area colocalized with Pavarotti. n=18-26 cells from 3 independent experiments. Ctrl = 23.5 ± 2.4%, Upper 95% CI = 28.6, Lower 95% CI = 18.3 Trc RNAi = 12.78 ± 1.9%, 16.62, 8.94. p = 0.001 Student’s T-test Scale bar = 10μm. G Example images of extracted S2 cells expressing mCherry Tubulin and WT Pavarotti GFP or Pavarotti S745A GFP. Endogenous Pavarotti was depleted with dsRNA targeting non coding regions. H Quantification of microtubule area colocalized with Pavarotti. n=12-17 cells from 3 independent experiments. Scale bar = 10μm. WT = 29.4 ± 5.0%, Upper 95% CI = 40.4, Lower 95% CI = 18.4, S745A = 7.79 ± 1.8%, 11.5, 4.08. p = 0.0001 Student’s T-test I. Western blot of S745 phospho-Pavarotti from HEK cell lysates shows an increase in this species with Trc overexpression.

Next, we sought to confirm that phosphorylation at S745 was necessary for Pavarotti to inhibit microtubule sliding. We once more carried out sliding assays, this time with a knockdown and rescue approach. Depletion of Pavarotti with a dsRNA against a non-coding region increased sliding which could be reduced again by expression of wild-type Pavarotti. However, expression of Pavarotti S745A mutant failed to reduce sliding levels to their control levels (Fig 3 C, D, video 6). Therefore, Pavarotti can no longer inhibit microtubule sliding when phosphorylation of S745 is prevented.

To understand why phosphorylation of Pavarotti is required for sliding inhibition, we compared microtubule binding of GFP-Pavarotti in S2 cells under control conditions, after Trc depletion, or after mutation of S745. In order to visualize microtubule-bound Pavarotti, we extracted S2 cells with Triton X-100 under conditions preserving microtubules (see Materials and Methods). We found that depletion of Trc led to around a 50% decrease in association of Pavarotti with microtubules (Fig 3 E, F). Parallel experiments with the phospho-null mutant S745A show the same effect (Fig 3 G, H). Therefore, Pavarotti in its unphosphorylated state associates less with microtubules. Under these conditions, microtubule sliding is permitted.

### Inhibition of Microtubule sliding by Pavarotti requires interaction with 14-3-3 proteins

Our data so far show a role for phospho-regulation of Pavarotti in microtubule sliding, beyond its canonical function in cytokinesis. We chose to continue investigating parallels between mitosis and interphase/post mitotic microtubule sliding, this time testing the involvement of 14-3-3 proteins. These proteins have been shown to form a complex with Pavarotti dependent upon phosphorylation at the identified S745 site (Douglas et al., 2010; Fesquet et al., 2015). This association influences microtubule bundling (Douglas et al., 2010). We chose to further probe this mechanism, both with regards to Pavarotti S745 phosphorylation by Trc and microtubule sliding. Initially we carried out co-immunoprecipitation experiments of exogenous GFP-Pavarotti from HEK-293 FT cells. We found a robust interaction between WT Pavarotti and endogenous 14-3-3 ξ. Mutation of S745 to alanine, mimicking a non-phosphorylated form of Pavarotti, abrogated the interaction. Co-expression of exogenous Trc, generating a greater pool of phosphorylated Pavarotti at S745 (Fig 4 A), increases the interaction between Pavarotti and 14-3-3 ξ. Therefore, we confirm that phosphorylation of Pavarotti, by Trc, is indeed functioning to recruit 14-3-3 proteins.

**Figure 4.**
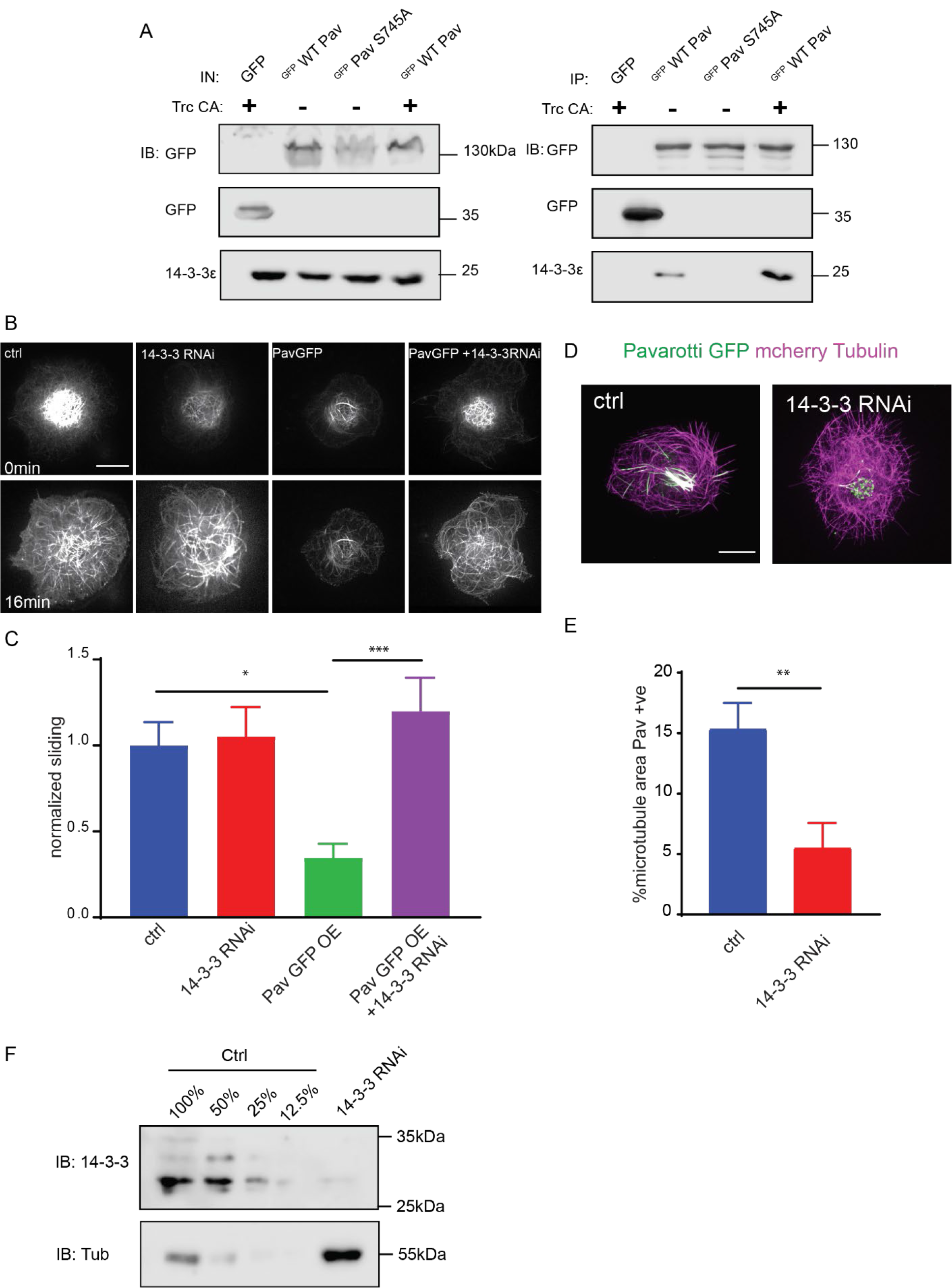
Phospho-Pavarotti brakes microtubule sliding via interaction with 14-3-3 proteins. A. Western blot from HEK cell lysate showing co immunoprecipitation of Pavarotti and 14-3-3. The interaction is lost upon mutation of S745 to Alanine and increased upon co-expression of the kinase Trc. B. Sliding experiments in S2 cells show the ability of Pavarotti to brake microtubule sliding is dependent upon 14-3-3 proteins as quantified in C. Scale bar = 10μm. n=21-26 cells from 3 independent experiments. Ctrl= 1 ± 0.14 Upper 95% CI = 1.28, Lower 95% CI = 0.72, PavOE= 0.35 ± 0.08, 0.52, 0.18 14-3-3 RNAi= 1.05 ± 0.17, 1.41, 0.70, 14-3-3RNAi + PavOE = 1.20 ± 0.20, 1.6, 0.8. ctrl vs pav OE p = 0.01, ctrl vs 1433RNAi p = 0.99, ctrl vs 1433 RNAi + PavOE p = 0.78, Pav OE vs 1433 RNAi p = 0.007, pav OE vs 1433 RNAi + Pav OE p = 0.0004, 1433 RNAi vs 1433 RNAi + Pav OE p = 0.91 One-way ANOVA with Tukey’s post hoc correction. D Example images of extracted S2 cells expressing mCherry Tubulin and WT Pavarotti GFP. Depletion of 14-3-3s decreases microtubule area decorated with Pavarotti. Scale bar = 10μm E. Quantification of microtubule area colocalized with Pavarotti. n=22-26 cells from 3 independent experiments. Ctrl = 15.4 ± 2.1, Upper 95% CI = 19.72, Lower 95% CI = 11.07, 14-3-3 RNAi = 5.54 ± 2.0, 9.78, 1.30. p = 0.0017 Student’s T-test. F. Western blot from S2 cell lysate demonstrating knockdown of 14-3-3.

We next tested if the interaction between Pavarotti and 14-3-3s is necessary for microtubule sliding inhibition by Pavarotti in S2 cells. We depleted *Drosophila* 14-3-3 β and ξ isoforms in S2 cells by dsRNA and overexpressed Pavarotti. Knockdown is demonstrated by western blot in Fig 4 F. Overexpression of Pavarotti caused a decrease in microtubule sliding. However, when we depleted 14-3-3 levels, we no longer observed this decrease in sliding upon Pavarotti overexpression (Fig 4 B, quantified in C, video 7). Thus, Pavarotti is not capable of inhibiting microtubule sliding in the absence of 14-3-3 proteins. These data are in good agreement with microtubule sliding assays performed with our S745A mutant, where a complex between Pavarotti and 14-3-3s does not form. Observation of the microtubule network in this condition showed a decrease in Pavarotti associated with microtubules (Fig 4 D, E), consistent with the effect seen with Trc depletion or the S745A mutation. Altogether, our data suggest Pavarotti locally brakes microtubule sliding by forming a complex with 14-3-3 proteins. The formation of this complex requires on phosphorylation at S745, by the kinase Trc.

### Pavarotti and Trc act in the same pathway to control dendrite outgrowth in vivo

We hypothesized that Trc and Pavarotti are in the same pathway in *Drosophila* neurons *in vivo* and regulate neurite outgrowth together. In order to test this, we measured the dendritic arbor of class IV DA (dendritic arborization) neurons (sensory neurons) in 3^rd^ instar larvae either upon knockdown of Trc, Pavarotti, or both in conjunction. Class IV neurons were labelled with ppk∷tdTomato and neuron specific expression of RNAis was achieved with elav gal4. In each case, we observed a roughly 40% increase in total dendrite length (Fig. 5 A, B). In the case of Trc, these data are consistent with previous reports (Emoto et al., 2004) where expression was abolished by mutation rather than neuron specific RNAi. Notably, the effect upon double knockdown was equivalent to that of the single knockdowns (Fig. 5 A, B). This supports the hypothesis that Trc and Pavarotti regulate neurite outgrowth in concert. Taken together, these data demonstrate the requirement for Pavarotti phosphorylation at S745 by Trc to brake microtubule sliding and correctly tailor neurite extension. Also, we show how Trc regulates microtubule sliding by influencing Pavarotti-microtubule binding.

**Figure 5.**
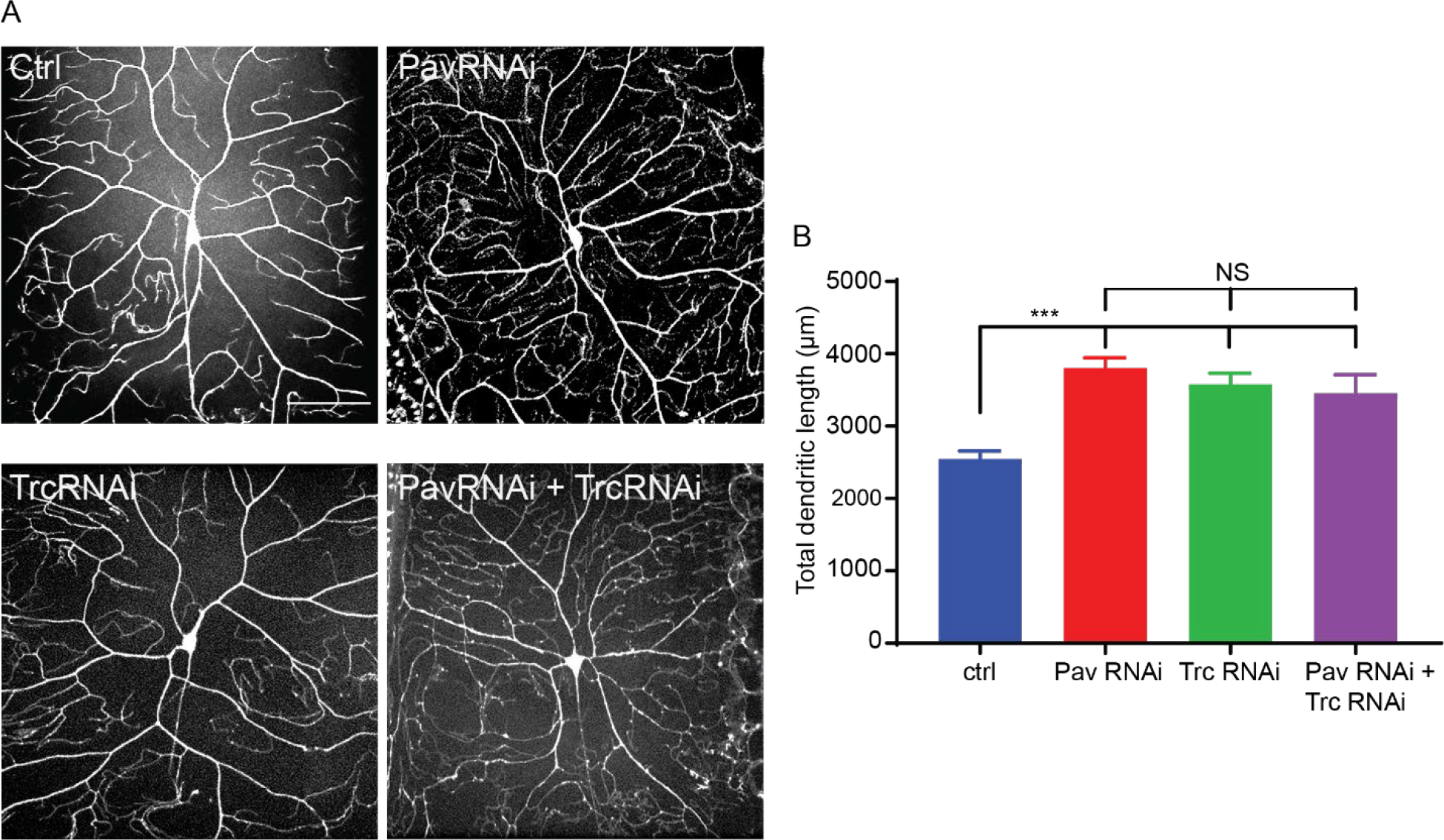
Pavarotti and Trc act in the same pathway to control dendrite outgrowth *in vivo*. A. Example images showing DA neurons labelled with ppk∷tdTomato in 3^rd^ instar larvae under control conditions, with Pavarotti or Trc RNAi driven by elavGal4, or both RNAis together. Scale bar = 50μm. n=15-30 cells from at least 4 animals. Ctrl = 2549 ± 108μm, Upper 95% CI = 2771, Lower 95% CI = 2328, Pav RNAi = 3803 ± 141μm, 4099, 3507, Trc RNAi = 3577 ± 155μm, 3904, 3250, Pav and Trc RNAi = 3455 ± 257μm, 4005, 2905. B. Pavarotti and Trc depletion cause an increase in dendritic length compared to control. Depletion of both proteins together has a non-additive effect compared with either RNAi. Ctrl vs Pav RNAi p < 0.0001, Ctrl vs Trc RNAi, p<0.0001, Ctrl vs double RNAi p = 0.0006, PavRNAi vs Trc RNAi p = 0.76, Pav RNAi vs double RNAi p = 0.48, Trc RNAi vs Double RNAi p = 0.96. One-way ANOVA with Tukey’s post hoc correction.

## Discussion

Developing neurons must extend neurites to form a network for correct communication. This outgrowth must be downregulated as the neurites reach their intended targets and form stable synapses. We have previously shown that microtubule-microtubule sliding is required for neurite outgrowth in young neurons and is diminished in mature neurons by the action of *Drosophila* kinesin-6, Pavarotti. However, the processes by which Pavarotti might temporally regulate microtubule sliding were unknown. Here we report an inhibitory pathway for microtubule sliding. Our previous work has demonstrated roles for “mitotic” processes (microtubule-microtubule sliding) and “mitotic” motors (kinesin-6 Pavarotti) in regulating the neuronal cytoskeleton. In this study, we extended these parallels to identify the NDR kinase Trc (Tricornered) as a required component of the Pavarotti pathway, regulating neurite outgrowth.

### Trc is a novel regulator of microtubule sliding

NDR kinases have well studied roles in cell division and tissue morphogenesis. The yeast homologue of Trc (Dbf2p) promotes chromosome segregation and mitotic exit. These functions are conserved in mammals (Hergovich et al., 2006; Tamaskovic et al., 2003). However, neuronal expression of some NDR kinases has additionally been reported. In neurons, depletion of Trc has been linked to increased outgrowth of both axons and dendrites across multiple taxa (Emoto et al., 2004; Gallegos and Bargmann, 2004; Ultanir et al., 2012; Zallen et al., 1999). Our data are in good agreement with these previous reports as we measure an increase in neurite length *in vitro* and an increase in dendrite length *in vivo*. Further, our data uncover a mechanism for this increased outgrowth. Depletion of Trc leads to increased microtubule sliding in both S2 cells and in cultured primary neurons. This increased sliding allows microtubules to push at the tips of nascent neurites, providing the force required for their extension (del Castillo et al., 2015; Lu et al., 2013). Indeed, this sliding has been shown to translate into dendrite outgrowth *in vivo* – expression of a sliding deficient kinesin-1 mutant drastically decreases the dendritic arbores of *Drosophila* sensory neurons (Class IV DA neurons) (Winding et al., 2016).

### Trc Phosphorylates Pavarotti to brake microtubule sliding

Similarly to Trc, the kinesin-6 Pavarotti was thoroughly studied with regard to cell division. It is a microtubule cross linker and signalling hub to promote cleavage furrow ingress (Adams et al., 1998; Basant and Glotzer, 2017; Verma and Maresca, 2019). Moreover, Pavarotti’s ability to localize to the spindle is dependent on its phosphorylation state (Guse et al., 2005). Here we have shown that Pavarotti is a downstream effector of Trc in the sliding inhibition pathway – Trc overexpression could only decrease sliding in the presence of Pavarotti. Using a similar approach, we have also demonstrated that Trc’s ability to regulate microtubule sliding is dependent upon its kinase activity. Knockdown and rescue experiments in S2 cells showed wild type and constitutively active Trc constructs could restore normal sliding levels but, a kinase dead variant was unable to do this. Further we have used a phospho-null mutant to demonstrate that phosphorylation of Pavarotti at the proposed Trc site of Serine 745 is necessary to inhibit sliding. Importantly, we confirm biochemically that *Drosophila* Pavarotti is phosphorylated at this site by Trc *in situ*. This extends previous reports to show this pathway is conserved between humans and *Drosophila* (Fesquet et al., 2015). *In vivo*, we demonstrate that these two proteins genetically interact to regulate neuronal development. Depletion of either Trc or Pavarotti leads to increased dendrite length in class IV DA neurons. Depletion of both of these proteins simultaneously has no additive effect, therefore these proteins act in a common pathway. Notably, this is the first report of Pavarotti regulating dendrite development in *Drosophila*. Previous work has shown Pavarotti prevents axon overgrowth (Del Castillo et al., 2015) and reports in mammalian systems have suggested roles in both compartments (Lin et al., 2012).

Whilst protein translation presents a clear alternative in regulating protein activity, we favour a phosphorylation model. Pavarotti expression is inhibited by Toll-6-FoxO signalling and Toll-6-FoxO mutants have increased microtubule stability (McLaughlin et al., 2016). However, phosphorylation would provide more dynamic method for modulating Pavarotti. Moreover, phosphorylation would provide tighter spatial regulation which could be necessary in inhibiting sliding in primary neurites while secondary processes are still developing. It is possible that Pavarotti phosphorylation inhibits kinesin-1 mediated microtubule sliding initially, and is subsequently regulated at the protein translational level.

### Phospho-Pavarotti forms a complex with 14-3-3 proteins to brake microtubule sliding

Extending our hypothesis that mitotic mechanisms regulating Pavarotti may be prevalent in neurons, we chose to investigate the role of 14-3-3s in microtubule sliding. 14-3-3s are conserved acidic proteins which bind phospho-threonine and phospho-serine residues (Cornell and Toyo-oka, 2017). Interaction and complex formation with phosphorylated proteins to facilitate cytoskeleton remodelling and axon extension has been described multiple times (Cornell and Toyo-oka, 2017; Taya et al., 2007). In *C. elegans* in mitosis, 14-3-3s have been shown to bind to the centralspindlin complex when Zen-4 (the *C. elegans* orthologue of Pavarotti/MKLP1) is phosphorylated at S710 (equivalent of Pavarotti S745) (Douglas et al., 2010). Here we show by co-immunoprecipitation experiments that Trc mediated phosphorylation at this site promotes formation of a complex between Pavarotti and 14-3-3s, in good agreement with previous data (Fesquet et al., 2015). This complex has previously been proposed to prevent stable microtubule binding *in vitro*. Interestingly, our data in S2 cells suggest the opposite. Examining the subcellular distribution of Pavarotti showed clear differences in microtubule association based on Trc-mediated phosphorylation state, and so, 14-3-3 interaction. We found Pavarotti localized more robustly to microtubules in the presence of Trc and that the phospho null mutant had a decreased ability to associate with microtubules. Upon association with microtubules, Pavarotti acts as a crosslinker and inhibits microtubule sliding. These observations are consistent with our sliding data – preventing Pavarotti phosphorylation by Trc upregulated microtubule sliding. Based on our findings, we suggest this phosphorylation promotes interaction with 14-3-3s, which in turn promotes microtubule localization. Indeed, our data show that 14-3-3s are necessary for Pavarotti to brake microtubule sliding.

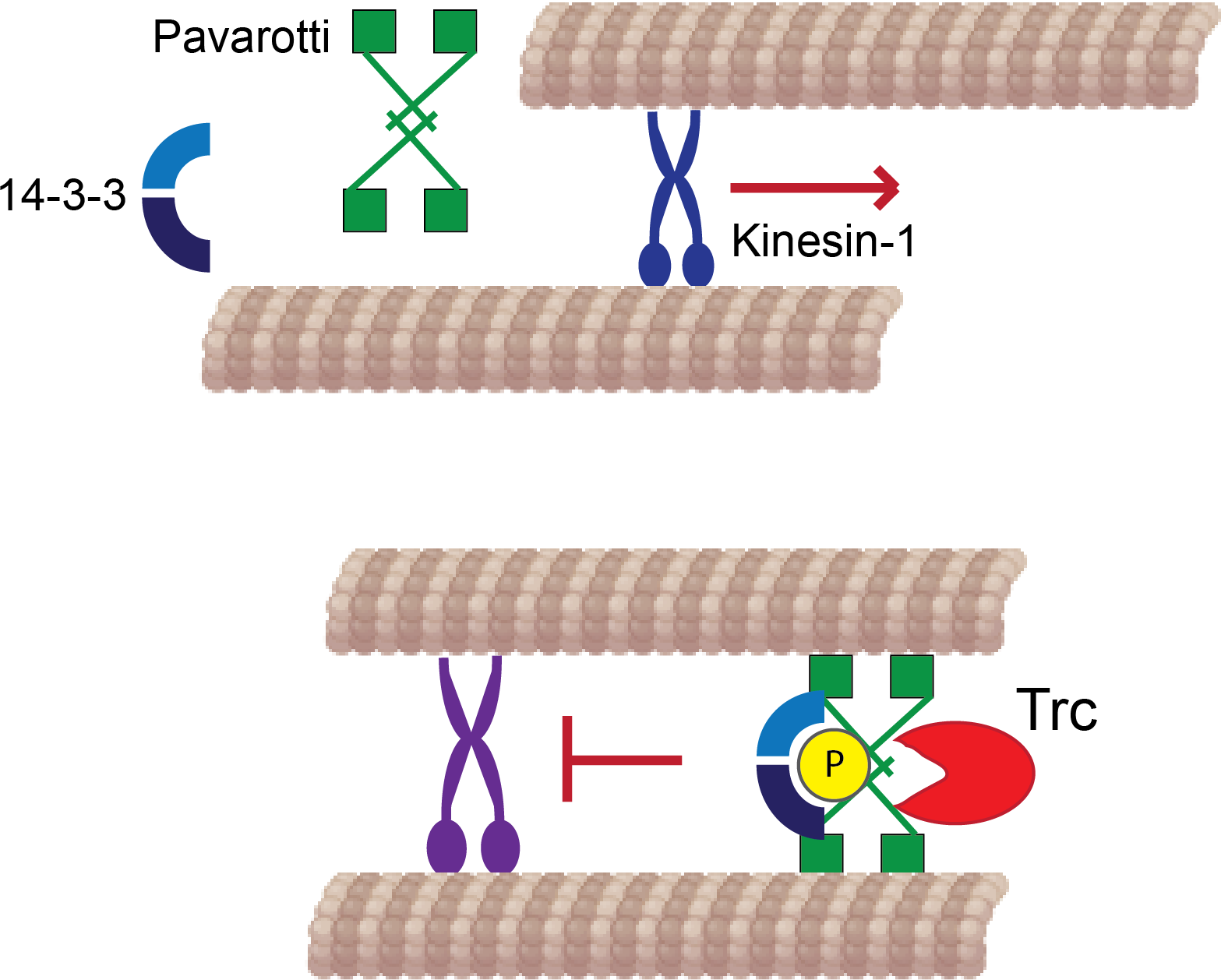
Summary of results. Kinesin-1 slides microtubules along one another to facilitate neurite outgrowth. The kinase Trc inhibits this process by phosphorylation of Pavarotti. Phosphorylated Pavarotti forms a complex with 14-3-3 proteins and associates with microtubules. Under these conditions, microtubules are cross-linked and can no longer undergo sliding by kinesin-1. Therefore, neurite outgrowth is inhibited.

Here we present a microtubule-based mechanism for Trc and Pavarotti controlling neurite development and show how the kinase Trc directs Pavarotti intracellular localization. Beyond this, it would be of interest to investigate a concurrent role for actin remodelling in process outgrowth. Pavarotti is a component of the centralspindlin complex (Mishima et al., 2002). The other component, tumbleweed/MgcRacGAP, gives an axon overextension phenotype upon depletion, like Pavarotti (Goldstein et al., 2005) and the expression levels of each are dependent on the other (Del Castillo et al., 2015). MgcRacGAP is a major orchestrator of RhoA signalling via Pebble/RhoGEF (Basant and Glotzer, 2018). This RhoGEF promotes formation of the cytokinetic furrow via actin assembly. Further, Trc has been proposed to inhibit Rac activity in a kinase dependent manner (Emoto et al., 2004). It has recently been demonstrated that the centralspindlin complex acts as a signalling hub to spatially and temporally regulate contraction of the actin cortex (Verma and Maresca, 2019). Might these signalling pathways be acting in developing neurites to tailor their development in addition to mechanical regulation via microtubule sliding?

The data presented here raise new questions beyond regulating microtubule sliding regulation. These data along with previous work from our group and others show a role for Pavarotti in controlling both axon and dendrite outgrowth (Del Castillo et al., 2015; Lin et al., 2012). This is consistent with our findings that kinesin-1 mediated microtubule sliding is necessary for proper development of both these compartments (Winding et al., 2016). Further, does microtubule sliding play a role in specifying axon formation? A crucial and distinctive feature of axons and dendrites is that of their microtubule polarity – axonal microtubules have a uniform, plus end out microtubule orientation, whereas dendritic microtubules are of mixed polarity or uniformly minus ends out (Baas et al., 1988; Stone et al., 2008). Kinesin-6 has been previously proposed to confer dendritic identity via transport of minus ends distal microtubules into dendrites and away from axons (Lin et al., 2012; Yu et al., 2000). Does the regulation of microtubule sliding via Pavarotti/kinesin-6 phosphorylation contribute to the microtubule polarity of nascent processes? How might these regulatory processes change over the course of axonal and dendritic development?

As well as neurite initiation, microtubule sliding occurs during axon regeneration (Lu et al., 2015). After axon or dendrite severing, *in vitro* or *in vivo*, large scale rearrangements of the microtubule cytoskeleton are observed (Lu et al., 2015; Stone et al., 2010). This is in contrast to in mature neurons where sliding is silenced. In this case, it could be of great interest to exploit our suggested mechanism of Pavarotti phosphorylation. Notably the kinase Trc may be a promising candidate for chemical inhibition. Would chemical inhibition or silencing of the kinase Trc deplete the pool of phospho-Pav and prolong the time period during which microtubule sliding was upregulated? Could this, in turn, facilitate neurite regeneration after injury? Further work will be required to address any potential for modulating Trc activity in neuronal regeneration.

## Materials and Methods

### Fly stocks

Flies were maintained at room temperature (24~25 C) on regular cornmeal food (Nutri-Fly, Bloomington Formulation), supplemented with dry active yeast. Stocks used in this study were: *w; elav-Gal4* (III, a kind gift from C. Doe), *yw; wg*^(*Sp*)^/*CyO; Dr*^(*Mio*)^/*TM3*, *Sb* (a kind gift from E. Ferguson), *yw; ppk-CD4-tdtomato* (II, BDSC stock 35844), *w; D42-Gal4* (III, BDSC stock 8816)(Pilling et al., 2006) *y sc v; UAS-Trc-RNA*i (TRiP.GL00028 and, TRiP.GL01127, attP2, BDSC stocks 35160 and 41591), *y sc v; UAS-Pav-RNA*i (TRiP.HMJ02232, attP40, BDSC stock 42573), *w; UASp-tdMaple3-alpha tubulin 84B* has been previously described (Lu et al., 2016). An insertion on the second chromosome was used to generate *yw; UASp-tdMaple3-alpha tubulin 84B; UASp-Trc RNAi* (TRiP.GL01127, attP2)

### Constructs and dsRNA generation

dsRNAs were generated using the sequences described below with the T7 sequence TAATACGACTCACTATAGGG at the 5’ end.

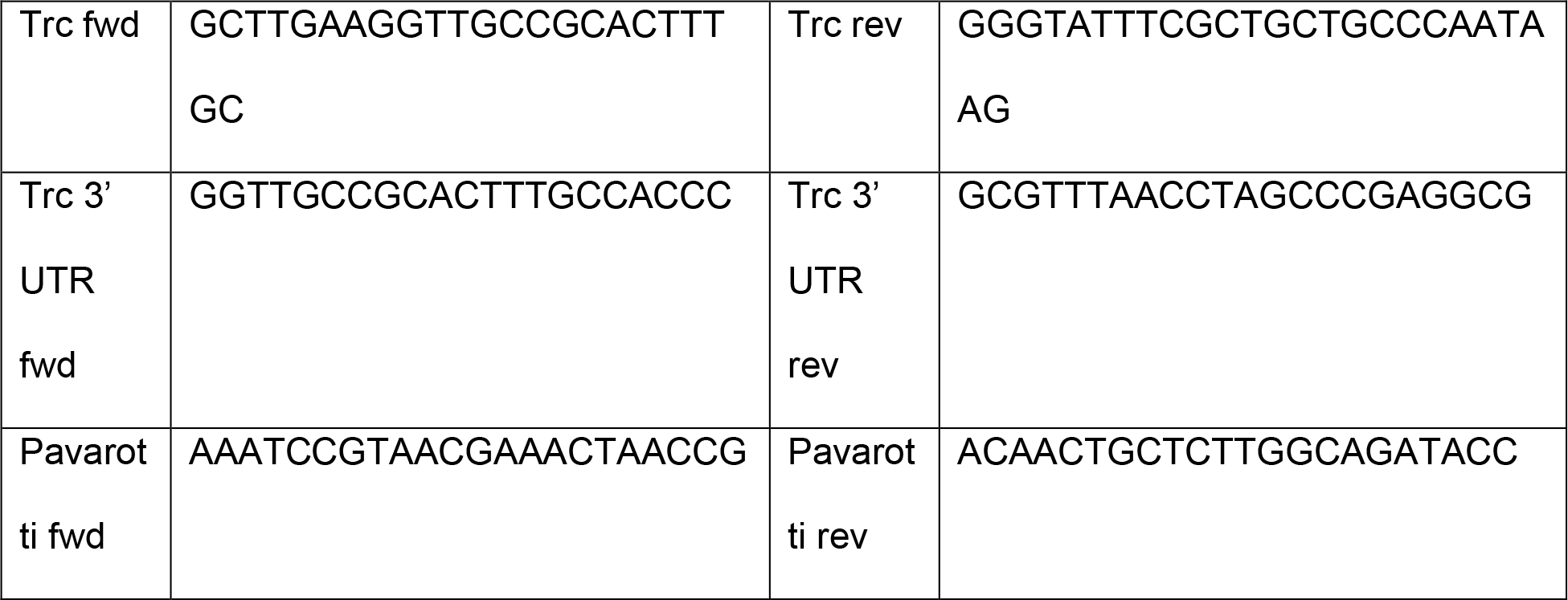
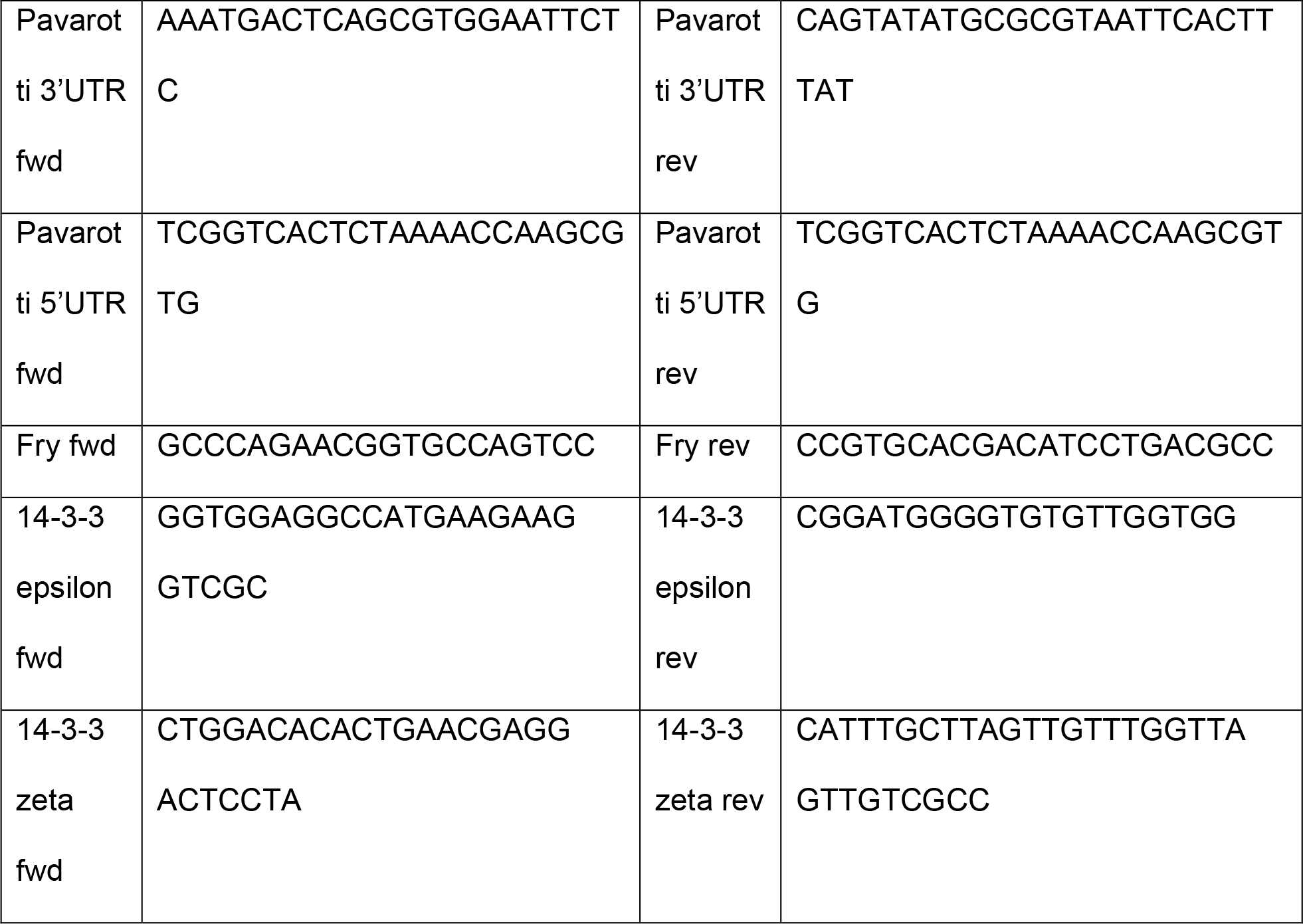

Constructs used in this study are: pMT EOS tubulin (described in (Barlan et al., 2013)). WT, T253E (constitutively active) and K122A (kinase dead) Trc constructs were a kind gift from P. Adler and were cloned from pUASt into pMT-BFP using EcoRI and NotI restriction sites. pMT GFP Pavarotti was generated from pMT-BFP Pavarotti, previously described in (Del Castillo et al., 2015). The phospho null mutant S745A was generated by site directed mutagenesis. For mammalian expression, Pavarotti constructs were subcloned in to pEGFP-C1 using EcoRI and SalI. Trc T253E was subcloned into pcDNA 3.1+ using HindIII and NotI. pMT-mcherry Tubulin has been previously described (Del Castillo et al., 2015).

### Cell culture

*Drosophila* S2 cells were maintained in Insect-Xpress medium (Lonza) at 25°C. Transfections were carried out with Effectene (Qiagen) according to the manufacturer’s instructions. dsRNA was added to cells on days 1 and 3 and imaging was carried out on day 5. HEK 293 FT cells were maintained in DMEM (Sigma Aldrich) supplemented with Penicillin Streptomycin and 10% FBS at 37°C and 5% CO_2_. HEK cells were transfected by Calcium Phosphate precipitation with 5μg DNA.

Primary neuronal cultures were prepared by dissection of brains from 3^rd^ instar larvae and dissociation of tissue using liberase (Roche). Cells were plated on ConA coated glass coverslips and maintained in Schneiders medium supplemented with 20% FBS, 5μg/ml insulin, 100μg/ml Pen-Strep, 50μg/ml Gentamycin and 10μg/ml Tetracycline. For sliding assays, larvae were cultured at 29°C and imaged 1hr after plating.

### Immunoprecipitation and western blotting

Co-Immunoprecipitation from HEK 293 cells was carried out in coIP buffer (50mM Tris pH 7.5, 150mM NaCl, 1.5% Triton X-100, 1mM EDTA, 1mM PMSF, 20μg/ml Chymostatin, Leupeptin, Pepstatin, 1mM NaVO_3_). For phosphorylation experiments, GFP-Pavarotti was enriched by GFP pull down in RIPA buffer (50 mM Tris pH 7.4, 150 mM NaCl, 1% Triton, 0.5% Na-Deoxycholate, 0.1% SDS, 1.5 mM NaVO_3_, 1 mM PMSF, 20μg/ml Chymostatin, Leupeptin, Pepstatin, 1mM NaVO_3_). Cells were lysed, debris was pelleted by centrifugation, and the soluble fraction was incubated with single chain anti GFP antibody (GFP-binder) (GFP-Trap-M; Chromotek) conjugated to sepharose beads. Samples were washed 3× in lysis buffer and boiled in 5× laemmli buffer prior to loading on 10% acrylamide gels for electrophoresis. To assess efficient knockdown of proteins, S2 cells were lysed directly in sample buffer and boiled. After electrophoresis, transfer onto nitrocellulose membrane was carried out and blocking was performed in 4% milk in PBS-T. For phospho- specific antibody, blocking was carried out with 3% BSA in TBS-T. Antibodies used were: Anti-Trc (a kind gift from K. Emoto), anti-Pavarotti (a kind gift from J. Scholey), anti-Tubulin DM1a, Anti-Pavarotti pS710 (corresponding to *Drosophila* S745) was a kind gift from M. Mishima, anti GFP was prepared in house. HRP conjugated mouse and rabbit secondary antibodies were from Jackson. Western blotting was performed using advansta western bright quantum substrate and Licor Imagequant system.

### Fixed imaging

For subcellular localization analysis of Pavarotti, S2 cells were plated on ConA coated coverslips and allowed to attach. Cells were then extracted in 30% glycerol, 1%triton, 1uM taxol in BRB80 for 3 minutes and imaged directly.

### Microscopy and photoconversion

To image dissociated neuronal cultures by phase contrast we used an inverted microscope (Eclipse U2000; Nikon Instruments) equipped with 60x/ 1.40-N.A objective and a CoolSnap ES CCD camera (Roper Scientific) and driven by Nikon Elements software.

To image *Drosophila* S2 cells and primary neurons, a Nikon Eclipse U200 inverted microscope with a Yokogawa CSU10 spinning disk confocal head, Nikon Perfect Focus system, and 100×/1.45-N.A. objective was used. Images were acquired with an Evolve EMCCD (Photometrics) using Nikon NIS-Elements software (AR 4.00.07 64-bit). S2 cells expressing tdEOS-tagged Tubulin were plated in Xpress with 2.5μM cytoD and 40nM taxol. For photoconversion of tdEOS-tagged Tubulin in sliding assays we applied 405-nm light from a light-emitting diode light source (89 North Heliophor) for 5s. The 405nm light was constrained to a small circle with an adjustable diaphragm, therefore only a region of interest within the cell was photoconverted. After photoconversion, images were collected every minute for >10 min.

### Microtubule sliding analysis

Analysis was carried out as previously described. Briefly, time-lapse movies of photoconverted microtubules were bleach-corrected and thresholded and the initial photoconverted zone was identified. The number of pixels corresponding to MTs was measured in total or outside the initial zone for each frame. The motile fraction (defined as MTs^outside_initial_zone^/MTs^total^) was plotted against time and the slope of the linear portion was calculated to represent microtubule sliding rate. For analysis of microtubule sliding in neurons, images were bleach corrected and denoised using despeckle in FIJI. Movies were then processed using the WEKA trainable segmenter in Fiji to generate probability maps of photoconverted microtubules (Arganda-Carreras et al., 2017). Probability maps were thresholded and analyzed in the same way as S2 cells.

### Statistical analysis and data presentation

Data are presented as mean ± standard error. Statistical analysis was carried out in GraphPad. Data were analyzed using student’s T-test or One-way ANOVA with Sidak’s post hoc correction for multiple comparisons. Data are collected from at least 3 replicates. Statistical significance is presented as * p<0.05, **p<0.01, ***p<0.001. Data in figure legends are presented as mean ± standard error, Upper 95% Confidence Interval, Lower 95% Confidence interval.

## Acknowledgements

This work was supported by NIH R01 GM052111 to V. Gelfand. We thank members of the Gelfand lab and M. Glotzer for helpful discussion. We thank the Bloomington Stock Center (NIH P40OD018537) for fly stocks. We thank M. Mishima for phospho specific Pavarotti antibodies, K. Emoto for Trc antibody and we thank P. Adler for Trc DNA constructs. We thank M. Winding for initial observations regarding 14-3-3 and microtubule sliding.

